# mRNA-decapping associated DcpS enzyme controls critical steps of neuronal development

**DOI:** 10.1101/2021.05.10.443481

**Authors:** Iva Salamon, Geeta Palsule, Xiaobing Luo, Alfonso Roque, Shawn Tucai, Ishan Khosla, Nicole Volk, Wendy Liu, Huijuan Cui, Valentina Dal Pozzo, Petronio Zalamea, Xinfu Jiao, Gabriella D’Arcangelo, Ronald P Hart, Mladen-Roko Rasin, Megerditch Kiledjian

## Abstract

Homozygous mutations in the gene encoding the scavenger mRNA-decapping enzyme, DcpS, have been shown to underlie developmental delay and intellectual disability. Intellectual disability is associated with both abnormal neocortical development and mRNA metabolism. However, the role of DcpS and its scavenger decapping activity in neuronal development is unknown. Here, we show that human neurons derived from patients with a DcpS mutation have compromised differentiation and neurite outgrowth. Moreover, in the developing mouse neocortex, DcpS is required for the radial migration, polarity, neurite outgrowth and identity of developing glutamatergic neurons. Collectively, these findings demonstrate that the scavenger mRNA decapping activity contributes to multiple pivotal roles in neural development, and further corroborate that mRNA metabolism and neocortical pathologies are associated with intellectual disability.

## Introduction

The control of gene expression during neuronal development is highly regulated at multiple levels including RNA turnover, which serves a critical role in establishing the steady state accumulation of mRNA in cells (DeBoer EM et al. 2013; Popovitchenko T and MR Rasin 2017). A key modulator of RNA stability is the removal of the protective N7-methylguanosine (m^7^G) cap at the 5’ end termed decapping. One class of decapping enzyme functions on the intact capped mRNA to remove the 5’ cap and initiate degradation of the mRNA from the 5’ end and include the Dcp2, Nudt3 and Nudt16 decapping enzymes (Grudzien-Nogalska E and M Kiledjian 2017). A second class consists of the scavenger decapping enzyme, DcpS, which functions to hydrolyze the resulting cap structure following decay of an mRNA from the 3’ end (Wang Z and M Kiledjian 2001; Liu H et al. 2002).

Mutations in DcpS can lead to varying motor and developmental delays, and cognitive defects (Ahmed I et al. 2015) ranging from mild to more severe phenotypes that also include microcephaly, musculoskeletal and craniofacial abnormalities referred to as Al-Raqad syndrome (ARS) (Ng CK et al. 2015; Alesi V et al. 2018; Masoudi M et al. 2020). This is an intellectual disability disorder identified in four families that harbor mutations in their *DcpS* gene that negatively impacts decapping activity. Despite the role of DcpS in cognition being well established, the mechanism by which DcpS exerts its function on brain activity remains elusive.

Extensive studies have shown a link between disrupted RNA metabolism, neuronal production and maturation, particularly during mammalian neocortical development with cognitive disorders, including intellectual disabilities and autism-spectrum disorders (Monuki ES and CA Walsh 2001; Rakic P 2009; Lui JH et al. 2011; Kwan KY et al. 2012; Popovitchenko T and MR Rasin 2017; Molnar Z et al. 2019; Kalebic N and WB Huttner 2020; Molnar Z et al. 2020). Fine-tuned spatiotemporal orchestration of cellular and molecular processes, including RNA metabolism that take place during prenatal brain development is a prerequisite for the proper generation of the remarkable neuronal diversity placed within the six functionally distinctive layers of the postnatal neocortex (DeBoer EM *et al.* 2013; Popovitchenko T and MR Rasin 2017; Zahr SK et al. 2019; Popovitchenko T et al. 2020; Hoye ML and DL Silver 2021). The neocortical neurogenic phase begins when neural progenitors called radial glial cells (RGC) start to give rise to spatiotemporally distinct glutamatergic neurons that can be traced back to their birthday (Rakic P 1988). RGC divide asymmetrically in the ventricular zone (VZ) to give rise to either postmitotic immature neurons or intermediate progenitor cells (IPC) which undergo further cell divisions in the germinative subventricular zone (SVZ) before generating more neurons.

All postmitotic immature neurons born either in the neocortical VZ or SVZ migrate radially to reach their layer destination in the cortical plate (CP). In an early step of radial migration, when post-mitotic neurons are starting their journey from the VZ into the SVZ, they will lose their bipolar morphology and acquire a multipolar shape, which is characterized by dynamic neurite extension and retraction and somewhat random movement in the SVZ. These multipolar migrating neurons subsequently undergo reversal of bi-polarity in order to resume their radial migration towards the CP using the RGC process (Rakic P 1972; Kriegstein AR and SC Noctor 2004). This multipolar-to-bipolar transition is a highly vulnerable stage that appears to be affected in a number of neurodevelopmental disorders (LoTurco JJ and J Bai 2006).

Once migrating neurons reach the CP, they will differentiate into distinct subpopulations of glutamatergic neurons with layer-specific identities (LoTurco JJ and J Bai 2006; Rakic P 2007; Lui JH *et al.* 2011; Popovitchenko T and MR Rasin 2017). The first waves of radially-migrating glutamatergic neurons settle in the deeper parts of the six-layered CP and differentiate into subcortically-projecting pyramidal neuronal subtypes. These lower layer (LL) neurons express specific markers, such as transcription factors Ctip2 (mostly layer V) and Tle4 (mostly layer VI). Later-born glutamatergic neurons bypass deeper layers and reach the appropriate superficial laminar positions (layers II-IV) of the developing CP (Rakic P 1974). Layer-specific transcription factors, such as Brn1 and Satb2, are responsible for the specification of these migratory neurons into intracortically-projecting neurons or upper-layer (UL) neurons that project ipsilaterally within the same hemisphere or contralaterally into the opposite hemisphere through the corpus callosum (Molyneaux BJ et al. 2007; Kwan KY *et al.* 2012).

Modulation at the level of RNA metabolism is emerging as a significant contributor to neocortical development and neuronal specification (DeBoer EM et al. 2014; Kraushar ML et al. 2014; Pilaz LJ and DL Silver 2015; Popovitchenko T et al. 2016; Zahr SK *et al.* 2019; Popovitchenko T *et al.* 2020; Hoye ML and DL Silver 2021). However, there is still a paucity of data regarding the contribution of distinct RNA processing defects to altered corticogenesis. In particular, the role of DcpS and mRNA decay overall during neocortical development remains poorly understood.

Here, we demonstrate the importance of an mRNA stability factor in the regulation of proper human neuronal and mouse neocortical development. We show that induced pluripotent stem cell (iPSC)-derived neurons from a patient with DcpS mutations exhibit decreased neuronal differentiation capacity and alterations in neurite elongation. Moreover, after silencing or overexpressing DcpS in a mouse model, DcpS is found to regulate the multipolar morphology of neurons during early radial migration and significantly contributes to neurite outgrowth and neuronal subtype specification. Together, our results provide new insight into the role of the DcpS protein during neuronal development, which is critical for understanding how regulation of mRNA stability contributes to normal neocortical development.

## Materials and Methods

### Neuronal differentiation of iPSCs

Lymphoblast cells from individuals IV-3 harboring homozygous DcpS mutation and the DcpS mutant heterozygote individual III-3 reported in (Ahmed I *et al.* 2015) were used to generate iPSCs. iPSCs were generated by RUCDR Infinite Biologics^®^ from human primary lymphocytes using Sendai viral vectors (CytoTune™, ThermoFisher Scientific), as previously described (Moore JC et al. 2012). iPSCs were maintained in mTeSR™ hPSC medium (STEMCELL Technologies). Cells were passaged either using ReLeSR™ passaging reagent (STEMCELL Technologies) or with Accutase and cultured onto Matrigel-coated six-well plates. Induced cortical neurons (iNs) were generated directly from human iPSCs using an inducible lentiviral system, as previously described (Zhang Y et al. 2013). Briefly, iPSCs were plated in the presence of the Y-compound (5μM) and transduced with lentiviruses encoding rtTA (FUW-M2rtTA) and *NGN2* (Tet-O-Ngn2-puro). Lentivirus vectors were generously provided by Dr. Zhiping Pang (Rutgers, RWJMS). The next day *NGN2* expression was induced by the addition of doxycycline (2 μg/ml) to Neurobasal culture medium containing B27 supplement (Invitrogen), and infected cells were selected for 48h using puromycin (2 μg/ml). Cells were cultured for additional 5-7 days in B27- and doxycycline-containing Neurobasal medium.

### iN immunofluorescence and high-content analysis of microscopy images

Undifferentiated (day 0) or differentiated (day 7-9) iNs were washed twice with cold 1xPBS, fixed with 4% paraformaldehyde, permeabilized with 0.4% Triton X-100, and incubated in blocking buffer containing 10% normal goat serum. Cells were incubated for 1h at room temperature with primary antibodies in a buffer containing 0.4% Triton X-100 and 10% normal goat serum. Primary antibodies were: HuC/D (ThermoFisher # A-21271; 1:250 dilution) and TuJ1 (BioLegend # MMS-435P; 1:250 dilution). Cells were washed in 1xPBS, incubated with secondary antibodies for 1h, and washed again in 1xPBS before staining with DAPI (ThermoFisher; 1:1000 dilution). Secondary antibodies were goat anti-mouse IgG conjugated to AlexaFluor-647 (CY5 red) or −488 (FITC green) (ThermoFisher). To determine the percentage of neuronal differentiation, HuC/D- or Tuj1-immunostained cells were imaged with the INCell Analyzer 6000 (GE Healthcare). The analysis between the control and the mutant cell lines was carried out at 20x magnification using images with comparable number of nuclei per image (from 3 independent experiments), and the averaged values of each image analyzed using the INCarta^™^ image analysis software (GE Healthcare). Statistical analysis and graphical representation of data was performed using the GraphPad Prism (version 8) software. Since all values were normally distributed, an unpaired *t*-test was used to compare subject groups.

### Reverse-Transcription (RT)-qPCR analysis of gene expression

RNA was extracted using TRIzol™ (Thermo Fisher Scientific) followed by isopropanol precipitation. Extracted RNA was treated with DNase I (Promega) and used for reverse transcription (Promega RT kit) to generate cDNA. Gene expression analysis was conducted using SYBR and the Applied Biosystems (Thermo Fisher Scientific) QuantStudio3 qPCR machine. The qPCR primers used for analysis are: Nanog P1: 5’-CTCCAACATCCTGAACCTCAG-3’, Nanog P2: 5’-sCGTCACACCATTGCTATTCTTCG-3’; Oct4 P1: 5’-CCTGAAGCAGAAGAGGATCACC-3’, Oct4 P2: 5’-AAAGCGGCAGATGGTCGTTTGG-3’; Sox2 P1: 5’-GCTACAGCATGATGCAGGACCA-3’, Sox2 P2: 5’-TCTGCGAGCTGGTCATGGAGTT-3’; Nestin P1: 5’-TCAAGATGTCCCTCAGCCTGGA-3’, Nestin P2: 5’-AAGCTGAGGGAAGTCTTGGAGC-3’; Pax6 P1: 5’-CTGAGGAATCAGAGAAGACAGGC-3’, Pax6 P2: 5’-ATGGAGCCAGATGTGAAGGAGG-3’.

### RNA-sequence (RNA-seq) analysis

Library preparation, quality control testing, and Illumina high-throughput RNA sequencing was performed by RUCDR, Rutgers University (Piscataway, NJ). Resulting fastq files were trimmed and filtered by quality score using Fastp (Chen S et al. 2018), and then aligned with human genome (hg38) using HISAT2 (Kim D et al. 2019). Deduplicated, sorted bam files were used to extract gene counts using the Rsubread package (Liao Y et al. 2019) in R/Bioconductor. Counts were modeled in DESeq2 (Love MI et al. 2014) and tested for differential gene expression using a Wald test with significance selected at 5% false discovery rate and 1.5-fold change. Functional analysis used the enrichr (Kuleshov MV et al. 2016), ggprofiler2 (Raudvere U et al. 2019) and rrvgo (Sayols S 2020) packages. Raw data was deposited to public domain with GEO accession number: GSE173603.

### Expression vectors

The DcpS-specific shRNA plasmid was purchased from Millipore-Sigma (DcpS MISSION shRNA; TRCN0000304847) and consists of the sequence: 5’-CTATCACCTGCACGTGCATTT-3’. Lentiviral particles were generated with pCMV-VSVG and psPAX2 packaging plasmids according to the manufacturer (Thermo Fisher Scientific). Virus-containing supernatant was collected 48 hours post transfection and used at a multiplicity of infection of 5. The MISSION TRC2 pLKO.5-puro (Millipore-Sigma) was generated similarly and used as a control.

The plasmid pJLM1-FLAG-hDcpS shRR containing a flag-tagged shRNA resistant human DcpS gene was generated in multiple steps. A PCR generated human DcpS open reading frame with forward primer 5’-aagcttggatccATGGCGGACGCAGCTCCTCAACTAG-3’ and reverse primer 5’-TCTATgcggccgcTCGAGtcagctttgctgagcc-3’ containing a BamHI and a NotI restriction enzyme sites respectively and inserted into the same site of pIREShyg-FLAG vector. The shRNA resistant (shRR) version of the plasmid was generated by mutagenesis PCR with forward primer 5’-GACATATATAGTACGTATCACTTGTTCCCTCC-3’ and reverse primer 5’-GTTTGAAAATTGTAATTGGAGCTCAGGG-3’, which was subsequently used as template for PCR amplification to add AgeI and EcoRI restriction sites with the following primers: forward primer 5’-gctagcgctaccggtcgccacc ATGGACTACAAGGACGACGATGACAAG-3’ and reverse primer 5’-TCTAGATTCGAATTCTGGATCAGTTATCTATGCGGCCGC-3’ and used to replace the EGFP cassette at the same sites in pJLM1-EGPP to generate the pJLM1-FLAG-hDcpS shRR plasmid used in the study. The Nestin-DcpS plasmid was generated by extracting the Flag-DcpS from pLJM1-FLAG-DcpS shRR plasmid with EcoRI digestion followed by Klenow fragment mediated blunt ending and subsequent digestion with AgeI. The released DcpS fragment was inserted into linearized pNestin-EGFP (Addgene; # 38777) vector between a blunt ended NotI site and AgeI sites.

The pCdk5r-Fezf2-IRES-GFP plasmid was a gift from Paula Arlotta (Harvard University). The plasmid pIRESpuro3-EGFP was constructed by excising the EGFP open reading frame from pEGFP-C1 vector (Clontech) with NheI and BamHI digestion and insertion into the same restriction sites of the pIRESpuro3 vector (Clontech) to generate pIRESpuro3-EGFP. The PCR derived human DcpS open reading frame flanked by BamHI and NotI from above was inserted into the same sites of pIRESpuro3-EGFP to generate pIRESpuro3-EGFP-hDcpS. To generate the Cdk5r-DcpS overexpression constructs under the Cdk5r promoter, the pCdk5r-Fezf2-IRES-GFP plasmid was digested with XhoI and HpaI, blunt ended and used as vector for insertion of the DcpS gene. The DcpS open reading frame was derived from the plasmid pIRESpuro3-EGFP-DcpS by XhoI, blunt ended by Klenow fragment, and further digested with XmnI and inserted into the above vector. Clones were confirmed by sequencing.

### Animals and *In utero* Electroporation (IUE) studies

All embryonic experiments were performed according to the ethical regulations of Rutgers University Medical School’s Institutional Animal Care and Use Committee. Adult pregnant female CD-1 mice used for timed pregnancies were purchased from Charles River Laboratories. The day of vaginal plug detection was considered embryonic day (E) 0.5. Gender of the embryos was not analyzed, and embryos of either gender were used in the experiments. Our IUE protocol has been described previously (DeBoer et al. 2014). Plasmids were IUE at E13 and embryos were allowed to live until E17. Coelectroporation was performed with approximately 1 μL of plasmid mix that consist of 4 μg/μL of the control vector, 4 μg/μL of the DcpS shRNAs or overexpression vectors, together with 4 μg/μl CAG-GFP reporter. The administered electroporation mix was delivered at 4:1 ratio (construct: reporter). For each IUE, at least two or three transfected neocortices were used in experiments as described in figure legends with the “n” value. Two images from each electroporated section were taken, and one to four sections of different transfected neocortices were used for cell quantifications.

### Primary neuronal cell cultures

Primary neuronal cell cultures were prepared as previously described (Popovitchenko at al. 2020). Briefly, cover slips were treated with 1M HCl, washed with 1x PBS and coated using 15 ug/ml poly-L-ornithine (Sigma-Aldrich #P3655) at room temperature. Poly-L-ornithine was replaced with 4 μg/ml of mouse laminin (GIBCO #1094706) overnight at 37 °C. E13 neocrtices were dissected in HBSS containing 0.5% D-Glucose (American Analytical #AB00715-00500) and 25 mM HEPES (GIBCO #15630-080) and dissociated as described (Popovitchenko et al. 2020). Dissociated cells were re-suspended in Neurobasal medium (GIBCO #1097077) containing Sodium Pyruvate (GIBCO #11360-070), 2 mM glutamax (GIBCO #35050-061), penicillin/streptomycin (Cellgro #30-001-C1) and B-27 supplement (GIBCO #17504). All cells were cultured at 37 °C with 5% CO2. Cultured cells were transfected using either Ctrl or *DcpS* shRNA lentiviral constructs. After 72 hours, RNA was isolated as described above using TRIzol and submitted for RNAseq.

### Immunohistochemistry (IHC) and Confocal Imaging

IHC on mouse brain sections was performed at E13, E15 and E17 as described previously in detail (Popovitchenko et al. 2020). Briefly, dissected embryonic brains at desired age were fixed in fixation buffer (pH 7.4) that consist of 4% paraformaldehyde (PFA, Sigma-Aldrich, # 158127, St. Louis, MO, USA) in phosphate buffered saline (1xPBS, Corning, # 21-040-CV, Manassas, VA, USA) for 8h at 4 °C. Three washes in 1× PBS for 5 min were performed to remove fixation solution, and mouse brains were placed in 30% sucrose to properly prepare them for later use. Next, brains were washed three times for 5 min in 1xPBS, embedded in 3% agarose (Lonza, # 50004, Rockland, ME, USA) and coronally sectioned (Leica VT1000S vibratome) at 70 μm. Free-floating sections were incubated in blocking solution with gentle shaking for 3h at room temperature. Blocking solution contains 5% normal donkey serum (Jackson ImmunoResearch, # 017-000-121, West Grove, PA, USA), albumin (Biomatik, # A2134, Cambridge, ON, Canada), 0.4%Triton (Sigma, X-100, St. Louis, MO, USA), 0.2% L-lysine (Sigma, # L5501, St. Louis, MO, USA), 0.2% Glycine (BDH, # BDH4156, Radnor, PA, USA). The sections were then placed in a primary antibody solution diluted in blocking solution with 0.4%Triton X-100 and incubated with gentle rocking for 16h at 4 °C. The day after, three washes for 5 min in 1x PBS were performed to remove primary antibody solution. Sections were then incubated in the secondary antibody solution which was resuspended in blocking solution without Triton X-100 for 2 h with gentle shaking at room temperature. Sections were again washed three times in 1x PBS and incubated in 1 μg/ml of DAPI (Fisher Scientific, # D1306, Waltham, MA, USA) for 10 min at room temperature. Lastly, sections were washed two more times for 5 min in 1x PBS and mounted with Vectashield mounting media (Vector Laboratories, # H1000, Burlingame, CA, USA). The following primary antibodies and dilutions were used: chicken anti-GFP (Aves, 1:1000, GFP-1020), rat anti-Ctip2 (Abcam, 1:250, ab18465), goat anti-Brn1 (Novus Biologicals, 1:600, NBP1-49872), goat anti-Tle4 (SCB, 1:150, sc-13377), mouse anti-Satb2 (Abcam, 1:250, ab51502), affinity purified rabbit anti-DcpS (Liu 2002) at 1:400 dilution. Appropriate species-specific Donkey secondary antibodies were purchased from Jackson ImmunoResearch and were used at 1:250 dilution. Mouse brain images were acquired with Olympus BX61WI confocal microscope using either 10× or 20× objectives and Fluoview FV-1000 was used for image processing by using the identical confocal settings. Brain images were merged and binned in software Gimp2.10.14.

### Cell specification analysis and reconstructions

Binning conditions were kept constant across an experiment. The rectangle with the width size of 200μm was drawn from (1) pia to (2) VZ and was split into ten equal sized bins. Imaging and cell counting were done in double blind fashion where neither the person imaging nor quantifying knew the experimental condition. Gfp+ alone or Gfp+ cells colocalized with a specific identity marker were quantified by using DAPI as a reference point to confirm cell nuclei. The percentage of colocalization between the layer markers of interest and Gfp+ cells was determined as the fraction of the total Gfp+ neurons. For the cell morphology analysis, images were opened in Fiji (NIH; ImageJ ver. 2.0.0-rc-69/1.52p) and all Gfp+ cells in the cortex were marked using the Cell Counter plugin (license GPLv3). First, 10 equal sized bins were created as previously described. Gfp+ cells were assigned one of the three morphologies: bipolar, multipolar or unspecified. The counted number of Gfp+ cells with particular morphology in each bin was determined as the percentage of the total Gfp+ cells.

For manual 3D reconstruction of dendrites and axonal arbors of the Gfp+ neurons per each experimental condition, multi-tiff Z-stack images were analyzed with off-site Neurolucida software in a double-blind fashion as described previously (Rasin MR et al. 2011). On average 14 neurons per section per each experimental condition were reconstructed. We have analyzed 80 *Gfp*+ neurons in control group and 217 *Gfp+* neurons in *Cdk5r-DcpS* OE group.

### Statistical analysis

The data were represented as graph bars with mean ±SEM by using the GraphPad Prism9 software unless otherwise noted. We performed Shapiro-Wilk normality tests to test check for normal distribution of the data. For pairwise comparisons we used either parametric unpaired t test with Welch’s correction or non-parametric Mann-Whitney test. For multiple comparisons, we used one-way ANOVA, followed by post-hoc Tukey’s multiple comparison test. Statistical tests and the number (*n*) of replicates were noted in the figure legends. The significance threshold was set to p < 0.05 and is reported as: *p < 0.05; **p < 0.01; ***p < 0.001; ****p < 0.0001.

## Results

### DcpS mutation disrupts human neuronal development *in vitro*

To deduce the molecular and cellular ideology underpinning the intellectual deficit in individuals with disrupted DcpS decapping activity, we generated induced pluripotent stem cells (iPSCs) using lymphoblast cells derived from individual IV-3 in (Ahmed I *et al.* 2015). This affected individual harbors a homozygous (c.636+1G>A mutation) and only produces the decapping deficient mutant DcpS protein with a 15 amino acid insertion after amino acid Q212 (DcpS-Mut). As a control, lymphoblast cells from a consanguineous cousin heterozygous for the G>A mutation (individual III-3 in (Ahmed I *et al.* 2015) was also used to generate iPSCs (DcpS-Ctrl) which produce both wild type and DcpS-Mut protein and are phenotypically normal (Ahmed I *et al.* 2015). The resulting DcpS-Mut and DcpS-Ctrl iPSCs were 94% and 96% double positive for the Oct4/Tra-1-60 pluripotency markers respectively (Figure S1A) and contained the expected elevated expression of Nanog, Oct4 and Sox2 mRNAs (Figure S1B).

The capacity of the DcpS scavenger decapping activity-deficient DcpS-Mut iPSCs to differentiate into neurons was next tested. Induced neurons (iNs) were generated by lentiviral transduction of the inducible *NGN2* gene to directly differentiate iPSCs into cortical excitatory neurons (Zhang Y *et al.* 2013). Following seven days of post induction, (D7iNs), a reduction of *Nanog, Oct4* and *Sox2* iPSC marker mRNA levels were observed along with an increase in *Nestin* mRNA (Figure S1B and S1C) consistent with neuronal differentiation. We next determined whether differential neurogenesis was evident between DcpS-Mut and DcpS-Ctrl cells by monitoring the expression of HuC/D, an RNA-binding protein specifically expressed in neurons (Chung S et al. 1996) and a marker for neuronal differentiation (Zucco AJ et al. 2018). Immunofluorescent detection of HuC/D and high-content analysis of microscopy images from D7-D9iN cells revealed a >3-fold increase in neuronal differentiation in the control cells relative to the DcpS mutant cells (Fig. 1A, B). Consistent with the reduced differentiation of the mutant DcpS into iN cells, Pax6 was higher in the mutant cells than in the control cells (Fig. 1C). These findings suggest attenuated neurogenesis in human cells harboring a decapping deficient DcpS protein.

**Figure 1.**
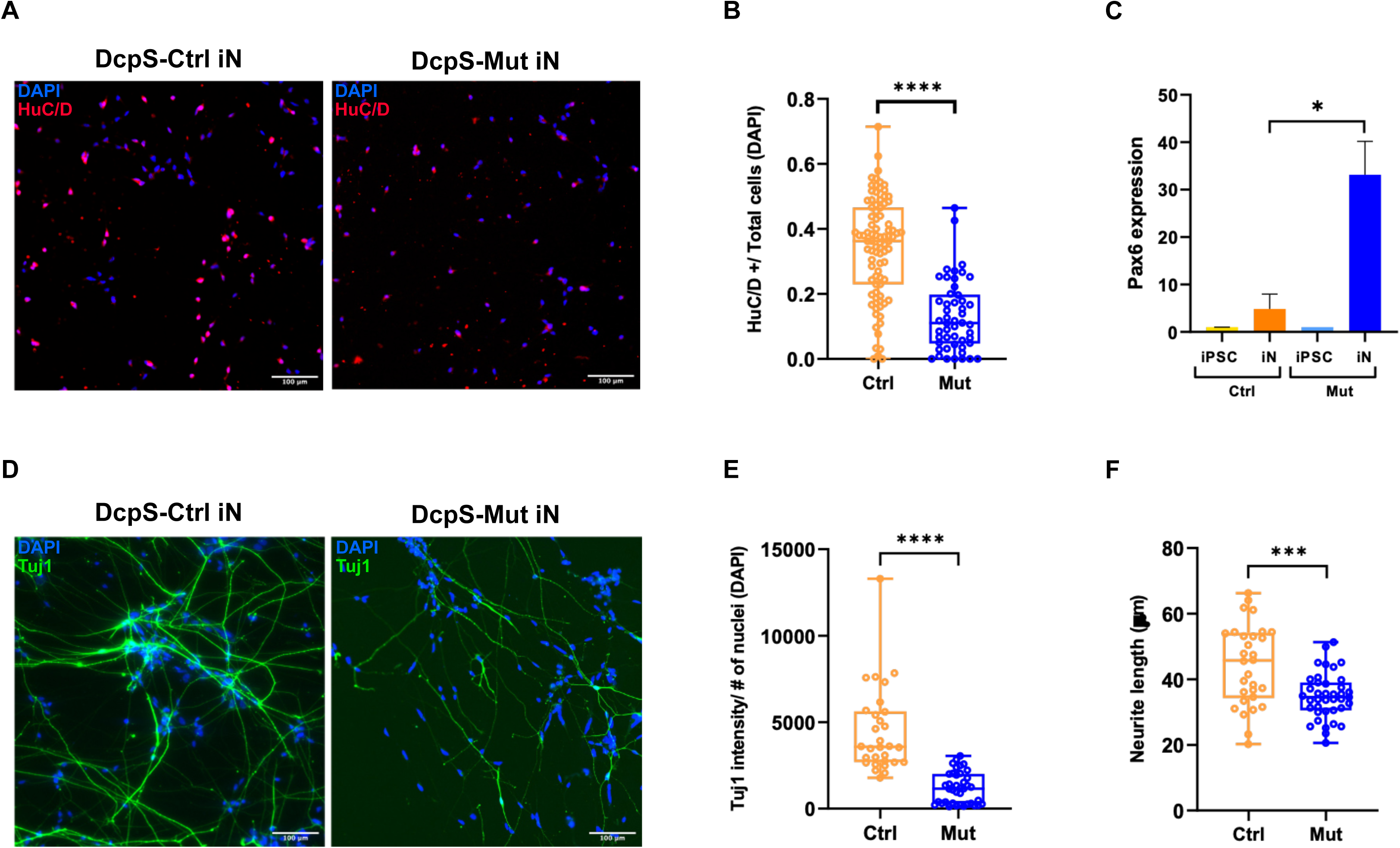
DcpS mutation disrupts human neuronal development *in vitro*. (**A**) Human induced Neurons (iNs) at Day 0 and 7-9 immunostained for neuronal marker HuC/D (red). DAPI stained nuclei are in blue. (**B**) Box and Whisker plot indicating the ratio of HuC/D positive cells at Day 7-9 to the DAPI positive cells per image. Images at the 20x magnification were collected and analyzed using the INCell Analyzer 6000 and the INCarta^™^ software. Statistics: DcpS-Ctrl, *n*= 89 images; DcpS-Mut, *n*=50 images from total 3 independent experiments; unpaired *t*-test, \*\*\*\**p*<.0001. (**C**) Pax6 expression detected by RT-qPCR, normalized to GAPDH expression. Statistics: *n*=3 biological replicates, 3 technical replicates for each; error bars indicate Standard error of mean (SEM); unpaired *t*-test, \*\**p*< 0.05. (**D**) Human iNs at Day 0 and 7-9 immunostained for Neuron-specific Class III β-tubulin (Tuj1) (green) to label neurites. DAPI positive nuclei are in blue. (**E**) Box and Whisker plot of the total Tuj1 intensity at Day 7-9 iNs normalized to the number of nuclei in each image. Images at the 20x magnification were analyzed using the INCarta™ image analysis software. Statistics: DcpS-Ctrl, *n*=30 images; DcpS-Mut, *n*=39 images from 3 independent experiments; unpaired *t*-test, \*\*\*\**p*<.0001. (**F**) Box and Whisker plot of the average neurite length (Tuj1 staining) per image for the DcpS-Het and the DcpS-Mut Day 7-9 iNs. Images at the 20x magnification were analyzed using the INCarta™ image analysis Software. Statistics: DcpS-Ctrl, *n*=30 images; DcpS-Mut, *n*=39 images from 3 independent experiments; unpaired *t*-test, \*\*\**p*<.001. Statistical analysis was performed using the GraphPad Prism 8 software. Error bars represent standard deviation (SD) unless otherwise stated.

The possible role of DcpS decapping in the outgrowth of neuritic projections was next analyzed. Automated image analysis of total immunofluorescence intensities of the neural-specific microtubule protein β-III-tubulin (Tuj1), revealed higher protein levels in D7-D9iN control compared to mutant cells (Fig. 1D and 1E), indicative of impaired neuritogenesis. Moreover, quantitation of aggregate neurite lengths demonstrated significantly greater length of projections in differentiated control cells relative to differentiated mutant cells (Fig. 1F) demonstrating more pronounced neuritogenesis in cells expressing wild type DcpS protein. Collectively, these findings indicate that neural differentiation and maturation is compromised in DcpS-Mut cells and suggest a role for the scavenger decapping activity in these processes.

### DcpS expression during mouse embryonic development

To define spatial expression pattern of DcpS in the mouse cerebral cortex, we first searched the transcriptome wide-gene expression atlas (www.genepaint.com). *In situ* hybridization analysis of mid-sagittal sections at embryonic stage E14.5 revealed the strongest DcpS signal in the forebrain, including the neocortex, thereby suggesting its role in neocortical development (Fig. 2A). To further confirm the distribution of DcpS expression in mouse neocortex, we used publicly available single cell (sc) RNA-seq analysis of E14.5 and postnatal (P)0 embryonic cortex (Loo L et al. 2019); http://zylkalab.org/datamousecortex). This scRNA-seq dataset showed DcpS expression in proliferating radial glial cells (classified as RG1/2/4) in the VZ and intermediate progenitor cells in the SVZ (classified as SVZ3) at E14.5, as well as the UL callosal and corticospinal projection neurons (classified as layer II/IV) in the P0 mouse neocortex (Fig. 2B). Using immunohistochemistry (IHC) analysis at E13, E15 and E17 of wild-type mouse neocortices (Fig. 2C), we found DcpS-positive (DcpS+) mitotic figures in VZ and SVZ at E13 and E15. To extend this observation to the LL cortical layers, we analyzed colocalization of DcpS and Ctip2, an established marker of LL subcortically projecting neurons. DcpS+ neurons were detected in CP between E13-E17 and were co-immunostained with Ctip2. These findings suggest that DcpS may play multiple roles in the developing neocortex, including neuronal development.

**Figure 2.**
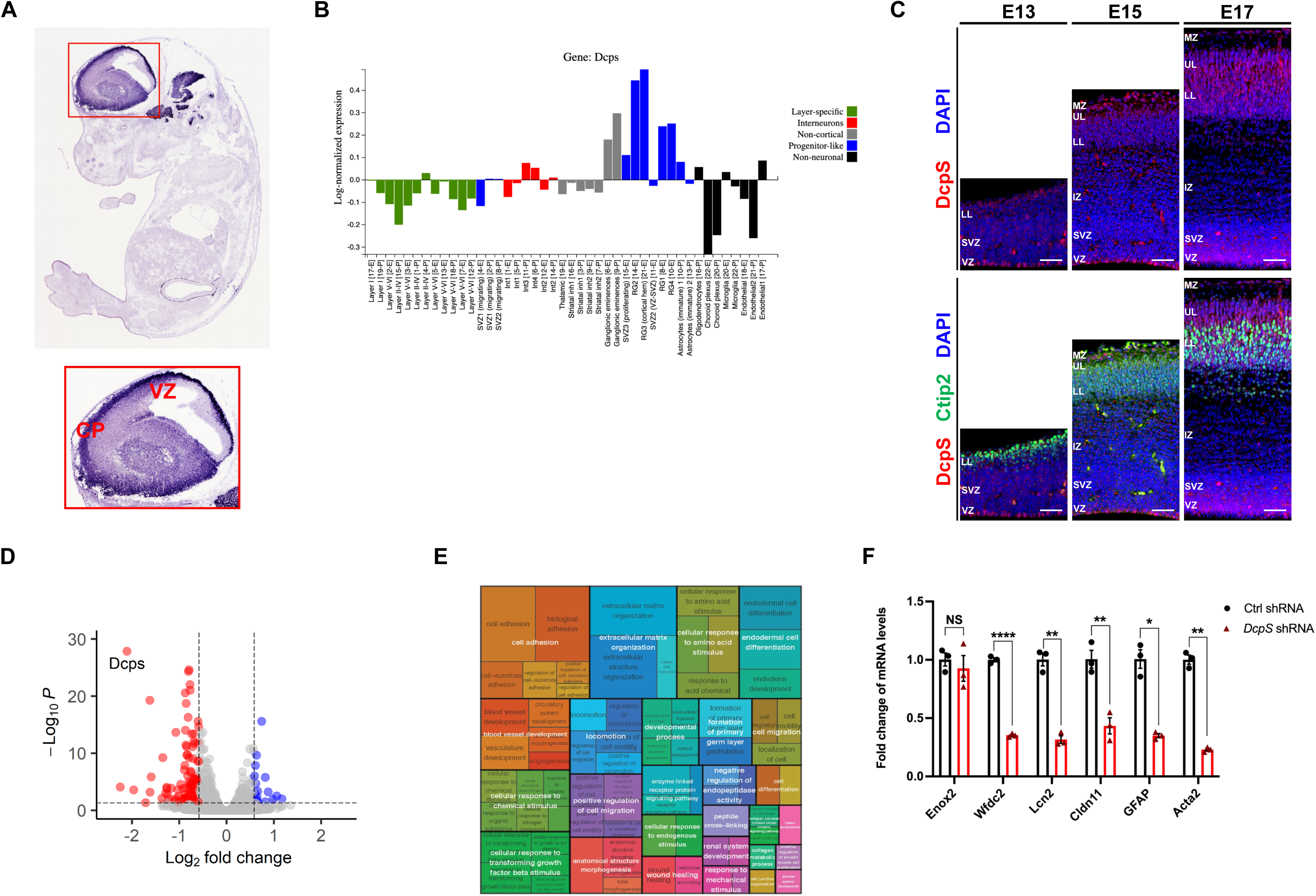
DcpS is expressed in developing mouse neocortex and regulates GO groups associated with neuronal development. (**A**) *DcpS in situ* hybridization at E14.5 from GenePaint (https://gp3.mpg.de). (**B**) Single cell RNA-seq analysis of E14.5 (E) and P0 (P) developing brain, including neocortex (Loo L *et al.* 2019); http://zylkalab.org/datamousecortex). (**C**) Representative confocal image of E13, E15, E17 immunostained for DcpS (red) and Ctip2 (green). DAPI is in blue. Scale bar of 20x objective lens: 50μm. VZ, Ventricular Zone. SVZ, Subventricular Zone. IZ, Intermediate Zone. LL, Lower-layer. UL, Upper-layer. MZ, Marginal Zone. (**D**) Volcano plot of transcripts altered by reduction of DcpS in primary mouse neurons. The log2 fold change (DCPS-KD/WT) is plotted versus the −log10 FDR (false discovery rate). Each transcript is indicated as a dot on the plot. Dots above the horizontal dashed line are <5% FDR. The vertical dashed lines indicate ±2-fold differences. Dots to the right of the +2-fold line are colored blue and to the left of the −2-fold are colored in red (only if they are detected with at least 1 FPKM). (**E**) REVIGO Treemap plot of GO Biological Process enriched terms generated from transcripts downregulated in DCPS-KD relative to WT. Individual terms are grouped by parent term within a colored region. Size of the rectangles reflect the p-value of enrichment, as calculated using the ggprofiler2 package in R. (**F**) qRT-PCR analysis of putative mRNAs affected by *DcpS* shRNA. *n*= 3 biological replicates, 2 technical replicates for each, error bars indicate Standard error of mean (SEM). Statistics: Welch’s t-test, unpaired, 2-tailed. *: p<0.05. **p < 0.01; ****p < 0.0001.

To explore the role of DcpS, we first used a DcpS knockdown (KD) approach to assess the effect of DcpS deficiency on global gene expression during neuronal development. Mouse primary neocortical neuronal cell cultures were transfected with either control (Ctrl) or *DcpS* shRNA. Unbiased RNA sequencing coupled to bioinformatics analysis identified 117 genes to be differentially expressed upon DcpS downregulation with 17 increasing and 100 decreasing (Fig. 2D). The *DcpS* mRNA was among the most reduced mRNAs with >4-fold reduction demonstrating the efficiency of the knockdown approach. Gene Ontology (GO) term analysis of the *DcpS* targets revealed that negatively affected genes in “biological processes” were enriched in “extracellular functions” including “extracellular matrix organization”, “cell adhesion” and “locomotion” (Fig, 2E). To validate the RNA-seq analysis, we tested mRNA levels of five target genes known to play a role in cell adhesion or extracellular matrix regulation by qRT-PCR: *Wfdc2, Lcn2, Cldn11, GFAP* and *Acta2.* The decrease in expression levels for each of these five target genes upon *DcpS* KD was highly comparable between the RNA-seq and qRT-PCR (Fig. 2F). As a negative control, the *Enox2* mRNA that was not influenced by *DcpS* KD in the RNA-seq was also not different by qRT-PCR. Collectively, our findings suggest that one potential class of genes that DcpS modulates may involve arranging and regulating the biological processes involved in neuronal development.

### Silencing and overexpression of DcpS disrupts radial migration and polarity of neocortical glutamatergic neurons

DcpS expression in developing neocortex and downstream targets associated with neuronal development processes, prompted us to investigate the role of DcpS *in* vivo. Unlike the relatively static environment of a cell culture plate, the maturation of a neuron is compartmentalized *in vivo* in the neocortex and undergoes several steps as described above. To examine whether DcpS is necessary for neuronal migration during mouse cortical development, we conducted KD experiments via *in utero* electroporation (IUE) of E13 mouse neocortices. After E13 IUE of the *CAG-GFP* construct with either Ctrl or *DcpS* shRNA into the VZ, we performed IHC analysis at E17 (Fig. 3A). The laminar distribution of neurons was assayed in 10 equally sized bins from the pial surface above the marginal zone (MZ; bin 1) to the apical edge of the VZ (bin 10). *DcpS* downregulation resulted in redistribution of *Gfp*-expressing (*Gfp*+) glutamatergic neurons in developing neocortex, suggesting that the radial migration of KD neurons was severely affected. While the Ctrl *Gfp*+ neurons entered the CP and positioned appropriately beneath the MZ by E17, *Gfp*+ neurons expressing *DcpS* shRNA exhibited migratory anomalies affecting their final intracortical positioning. The percentage of *Gfp*+ neurons was significantly decreased in most superficial CP bins representing UL/MZ in *DcpS* shRNA when compared to the Ctrl cortices (Fig. 3B). This difference was compensated by the significantly increased number of *DcpS* KD *Gfp+* neurons in lower CP bins representing LL (Fig. 3B). This finding suggests that *DcpS* silencing impairs the final steps of radial migration of glutamatergic neurons into the CP layers when UL are surpassing the LL and are at the stage of detachment from the RGC process and need to terminally translocate once the final position within the CP is reached {Gongidi, 2004 #7708}.

**Figure 3.**
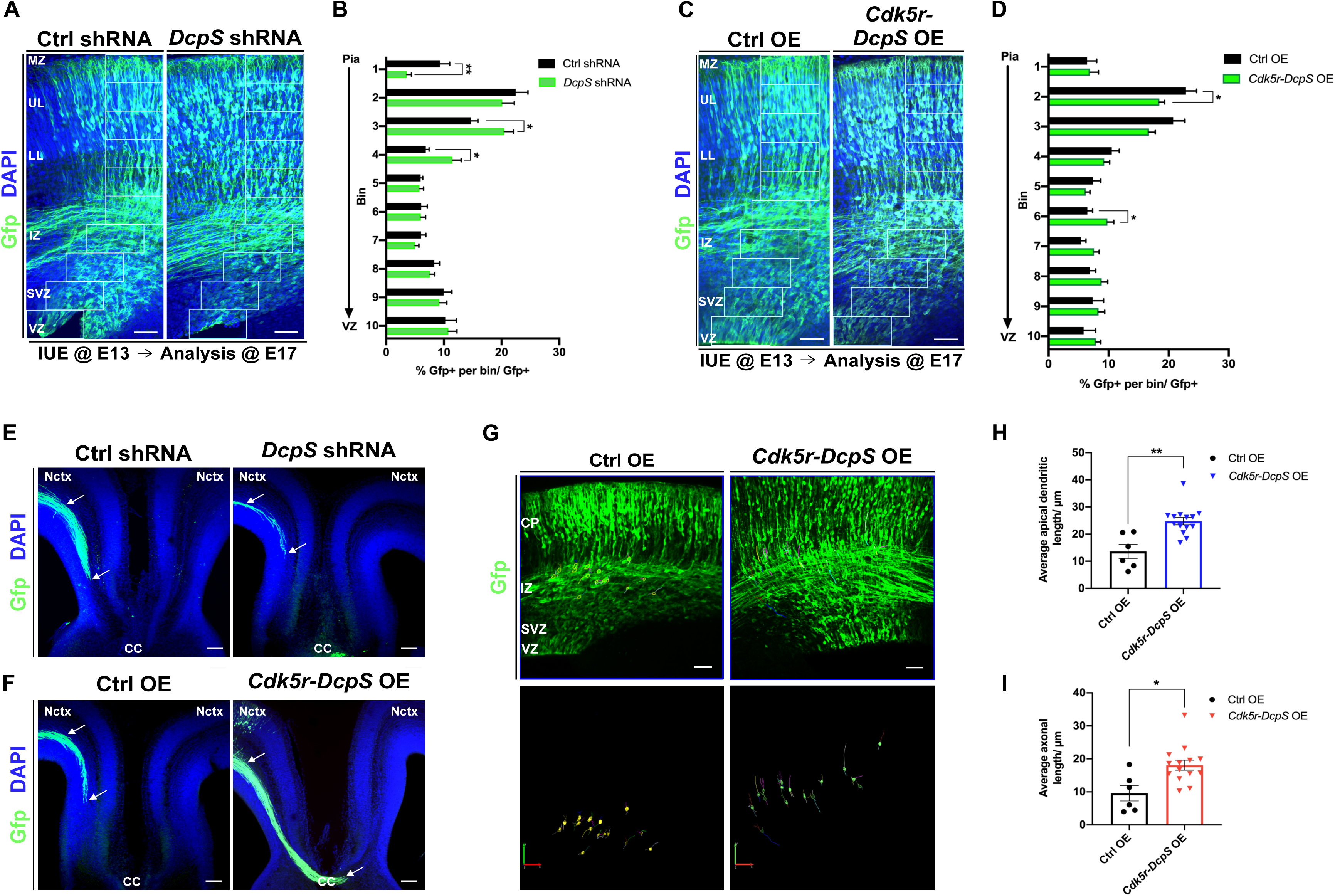
DcpS regulates radial migration of glutamatergic neocortical neurons. (**A**) Representative confocal images of E17 neocortices transfected at E13 with *Gfp* and either Ctrl or *DcpS* shRNA. DAPI is in blue. Scale bar of 20x objective lens: 50μm. (**B**) Quantification of distribution of *Gfp*+ cells in 10 equal bins from pial surface to VZ from A. *n*=18 images from 2 animals for Ctrl shRNA*; n*=35 images from 3 animals for *DcpS* shRNA. (**C**) Representative coronal confocal images of E17 neocortices transfected at E13 with *CAG-GFP* and Ctrl OE and *Cdk5r-DcpS* OE immunostained for *Gfp* (green) and DAPI (blue). Scale bar of 20x objective lens: 50μm. (**D**) Relative distribution of *Gfp+* cells subdivided in 10 equal bins for neuronal quantification from pial surface (bin 1) to VZ (bin 10) from C or E, respectively. Results represent *Gfp*+ neurons in each bin as the percentage from total *Gfp*+ neurons. *n*=12 images from 2 animals for Ctrl OE, *n*=29 images from 3 animals for *Cdk5r-DcpS* OE conditions. Statistics: Welch’s t-test per each bin, unpaired, 2-tailed. *: p<0.05. (**E**) Representative coronal images of corpus callosum (CC). Arrows denote *Gfp*+ axons arriving towards midline from transfected *Gfp*+ cells in in E17 Ctrl, *DcpS* shRNA neocortices or (**F**) E17 Ctrl OE and *Cdk5r-DcpS* OE neocortices. Scale bar of 10x objective lens: 100μm. (**G**) Representative images illustrating morphological threedimensional reconstructions of *Gfp*+ neurons migrating through IZ using Neurolucida from F. Scale bar of 20x objective lens: 50μm. (**H**) Quantitative analysis of apical dendritic lengths and (**I**) axon lengths in micrometers (μm) from H. *n*=80 neurons from 2 animals for Ctrl OE, *n*=217 neurons from 3 animals for *Cdk5r-DcpS* OE. Data represent the mean of sections and SEM. Statistics: Welch’s t-test per each bin, unpaired, 2-tailed, relative to the control. *: p<0.05, **: p<0.01. VZ, Ventricular Zone. SVZ, Subventricular Zone. IZ, Intermediate Zone. CP, Cortical Plate. LL, Lower-layer. UL, Upper-layer. MZ, Marginal Zone. CC, Corpus callosum. Nctx, neocortex.

To further assess the effect of DcpS on radial migration, DcpS was cloned under the control of the *Nestin* promoter, which is specifically restricted to mitotic RGC, or under the *Cdk5r* promoter, known to be active in early post-mitotic neurons (Rouaux C and P Arlotta 2013). E13 developing neocortices were IUE with *CAG-GFP* and either Ctrl overexpression (OE), *Nestin-DcpS* OE, or *Cdk5r-DcpS* OE constructs. At E17, we observed that *Gfp+* electroporated neurons reached the CP in Ctrl and cortices overexpressing *DcpS* under the control of the *Cdk5r* promoter (Fig. 3C and 3D). However, the proportion of *Gfp+* neurons was significantly decreased in the UL of *Cdk5r-DcpS* OE when compared to the Ctrl OE cortices. The observed difference was compensated by the significantly increased number of *Gfp+* neurons in the IZ of *Cdk5r-DcpS* OE. In contrast, *Nestin*-*DcpS* OE did not impair the radial migration as the distribution pattern of *Gfp*+ glutamatergic neurons remained unchanged when compared to the Ctrl OE (Fig. S2A and B). Our findings suggest that *DcpS* OE in progenitor cells does not interfere with the ability of their neuronal progeny to migrate. However, *DcpS* OE in postmitotic neurons decelerates their layer-specific radial migration into the CP.

Axonal and neurite growth is regulated by RNA metabolism (Hengst U and SR Jaffrey 2007; Batista AF and U Hengst 2016; Dalla Costa I et al. 2021). We next sought to determine whether E13 IUE *DcpS* KD or OE will influence the axonal growth of glutamatergic cortical neurons at E17. We found a reduction in the number of Gfp+ axons that reached the midline upon *DcpS* KD (Fig. 3E). In contrast, *DcpS* OE from the *Cdk5r* promoter favored the axon growth that overextended through the CC midline (Fig. 3F). To further assess if *DcpS* OE under the *Cdk5r* promoter induces changes in neurite extension, we reconstructed *Gfp*+ migrating neurons to analyze their complexity in detail (Fig. 3G). Sholl analysis showed that the average extension of both apically extending dendrites and axons increased in the *Cdk5r-DcpS* OE condition when compared to Ctrl OE (Fig. 3H, 3I). Together, our results indicate that DcpS promotes extension of leading processes in migrating glutamatergic neurons.

### DcpS is required for determination of multipolar morphology in post-mitotic neurons

Collectively, our findings suggest that DcpS may affect the polarity of neurons during neocortical migration as neurons undergo dynamic morphological transitions within the lower IZ and SVZ. Any perturbations in the transition from multipolar to bipolar morphology can lead to disruptions in radial migration. Thus, we investigated whether changes in DcpS expression levels affect their migration morphologies by quantitative analysis of *Gfp*+ neurons with acquired bipolar or multipolar morphology after E13-E17 IUE of *CAG-GFP* and either Ctrl OE or *Cdk5r-DcpS* OE (Fig. 4A). Distribution analysis of multipolar *Gfp+* neurons in *Cdk5r-DcpS* OE cortices showed a significant decrease in the LL accompanied by a modest increase in the lower IZ and SVZ (Fig. 4B). There was no apparent difference in the distribution of bipolar *Gfp*+ neurons when comparing *Cdk5r-DcpS* OE and control cortices (Fig. S3A).

**Figure 4.**
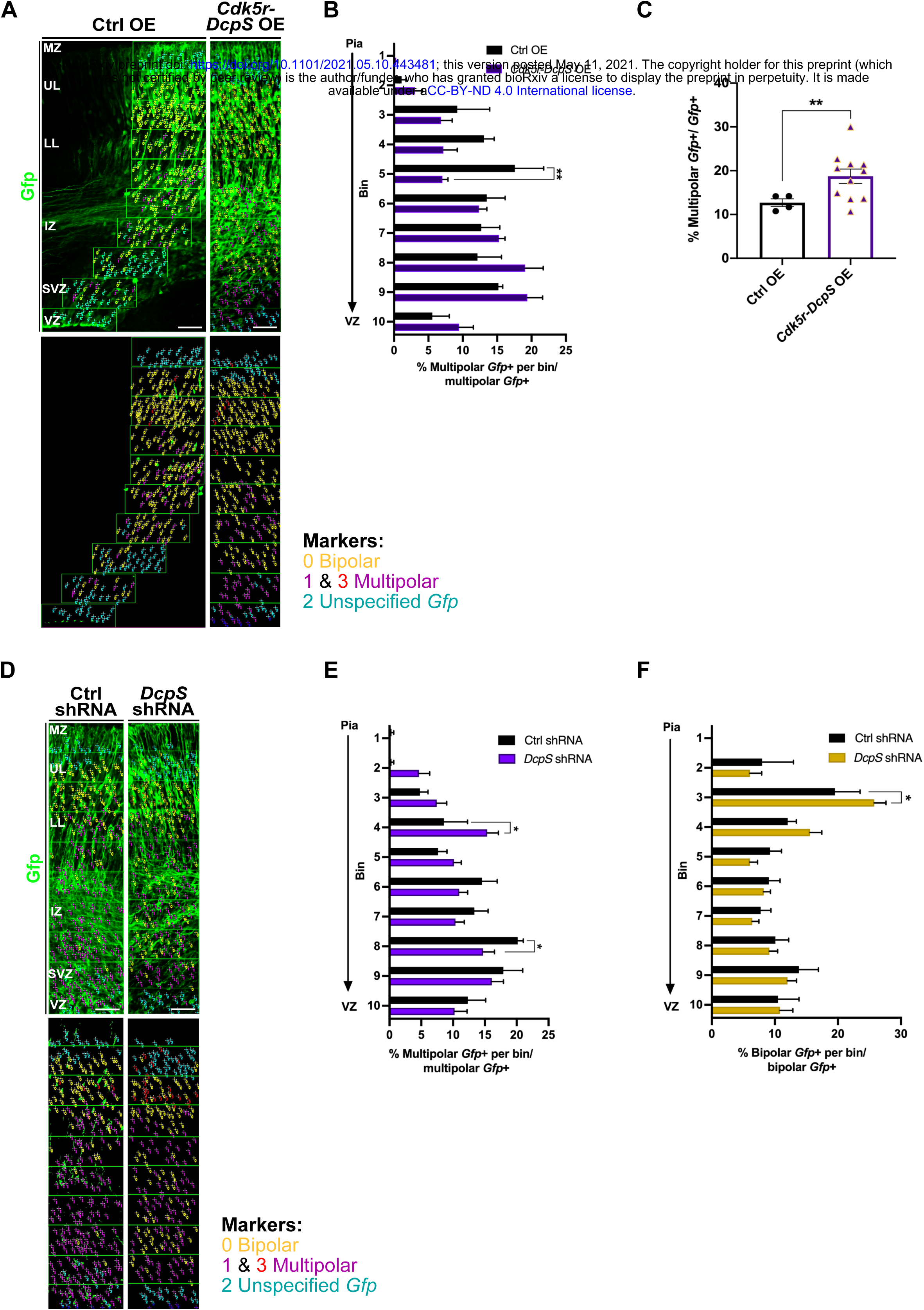
DcpS is required for the multipolar layout of neurons during radial migration. (**A**) Representative confocal images (top) and reconstructions of laminar positioning (bottom) of morphologically diverse *Gfp+* neurons of E17 neocortices transfected with Ctrl OE or *Cdk5r-DcpS* OE together with *CAG-GFP*. Scale bar of 20x objective lens: 50μm. (**B**) Quantified distribution of subpopulation of multipolar *Gfp+* neurons from A. Values are presented as percentage of morphologically specific *Gfp*+ neurons in each bin over the total morphologically specific *Gfp*+ neurons. (**C**) Quantification of the total proportion of multipolar *Gfp*+ neurons transfected with Ctrl or *Cdk5r-DcpS* OE from the total *Gfp*+ labeled cells. *n=* 4 images from 2 animals for Ctrl OE, *n*= 11 images from 3 animals for *Cdk5r-DcpS* OE condition. Data represent the mean of sections and SEM. Statistics: Welch’s t-test per each bin, unpaired, 2-tailed, relative to the control. *: p<0.05, **: p<0.01. (**D**) Representative confocal images (top) and reconstructions of laminar positioning (bottom) of morphologically diverse *Gfp*+ neurons of E17 neocortices transfected with Ctrl or *DcpS* shRNA with *CAG-GFP*. Scale bar of 20x objective lens: 50μm. (**E**) Quantification of the relative distribution of multipolar or (**F**) bipolar *Gfp+* cells per each bin is given as the percentage of the entire multipolar or bipolar *Gfp*+ cells, respectively, from D. *n*= 6 images from 2 animals for Ctrl shRNA, *n*= 14 images from 3 animals for DcpS shRNA. Data represent the mean of sections and SEM. Statistics: unpaired Welch’s t-test or Mann-Whitney test. NS= not significant, *: p<0.05. VZ, Ventricular Zone. SVZ, Subventricular Zone. IZ, Intermediate Zone. LL, Lower-layer. UL, Upper-layer. MZ, Marginal Zone.

We next assessed if DcpS might affect the balance of migrating neurons between multipolar and bipolar morphology. *Cdk5r-DcpS* OE cortices displayed a significant increase in the percentage of multipolar *Gfp*+ neurons (Fig. 4C). Interestingly however, a corresponding decrease in the total percentage of bipolar *Gfp+* neurons was not observed (Fig. S3B). These findings suggest that DcpS acts on retaining migratory neurons in multipolar stage but neither production of bipolar migratory neurons nor transition from bipolar to multipolar stage is affected.

To further examine the contribution of *DcpS* to neuronal morphology in the developing neocortex, we analyzed neocortices that were IUEd with *CAG-GFP* and either Ctrl or *DcpS* shRNA at E13 (Fig. 4D). At E17, *Gfp+* multipolar neurons in DcpS KD cortices were significantly increased in the lower CP but decreased in the lower IZ (Fig. 4E). Bipolar *Gfp*+ neurons had mild alterations in laminar distribution relative to the control, with sole increase at the borderline of UL and LL (Fig. 4F), whereas the total percentages of *DcpS* silenced bipolar and multipolar *Gfp*+ neurons were comparable to the Ctrl (Fig. S3C and S3D). Taken together, these data suggest that *DcpS* affects the migratory morphologies and localization of multipolar glutamatergic neurons during neuronal migration.

### Scavenger DcpS protein influences the identity of UL and LL glutamatergic neurons

Next, we assessed if *DcpS* KD affects the specification and/or laminar distribution of cortical glutamatergic neuronal identities at E17. IHC of specific neuronal subtypes was carried out with markers for UL and LL intracortically projecting (Satb2) and UL intracortically projecting (Brn1) neurons. *DcpS* silencing disturbed placement of Satb2+/Gfp+ post-mitotic neurons (Fig. 5A) with significantly reduced percentage of neurons in bins 1 and 7 (Fig. 5C). *DcpS* silencing also redistributed Brn1+ neurons when compared to the Ctrl (Fig. 5B). Significantly higher percentage of Brn1+/Gfp+ cells was localized at the border of UL and LL, while significantly lower percentages were detected surrounding these bins (Fig. 5D). Moreover, this altered distribution is associated with a significant reduction in the total percentage of Satb2+/Gfp+ and Brn1+/Gfp+ neurons (Fig. 5E, 5F), upon DcpS silencing. These findings further suggest that DcpS regulates last steps of UL migration.

**Figure 5.**
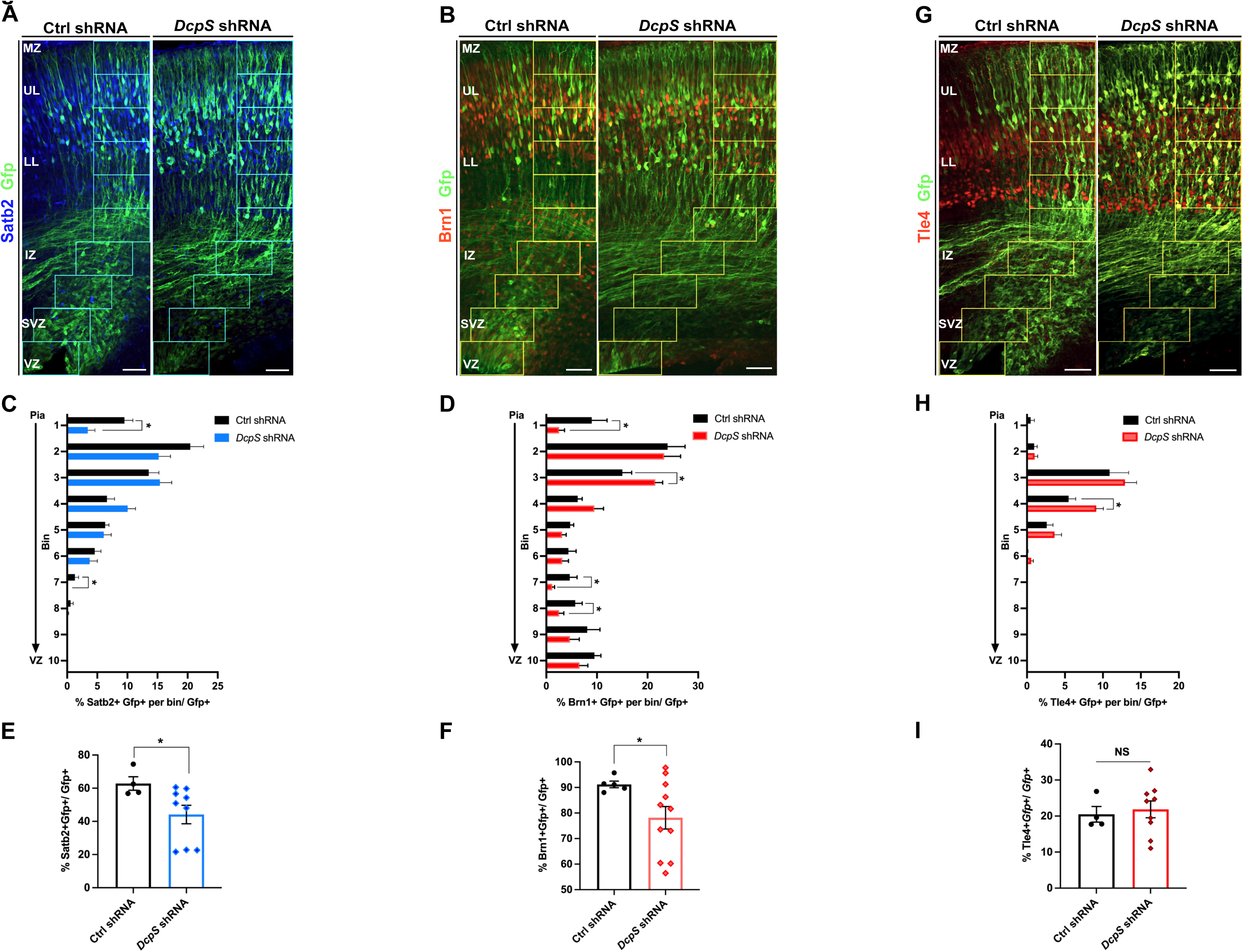
DcpS knockdown disrupts identities of upper layer neocortical glutamatergic neurons. Representative confocal images of E17 neocortices transfected at E13 with *CAG-GFP* and Ctrl shRNA or *DcpS* shRNA. Neocortices were immunostained for UL identity markers (**A**) Satb2 in blue and (**B**) Brn1 in red. Scale bar of 20x objective lens: 50μm. (**C**) Quantification of the laminar distribution of Satb2+ and (**D**) Brn1+ neurons from pial surface (bin 1) to VZ (bin 10), given as a percentage of the total number of *Gfp+* cells from A and B. (**E**) Quantification of the total number of identified *Gfp+* Satb2+ or (**F**) Brn1+ *Gfp+* neurons given as a percentage from total *Gfp+* cells. (**G**) Coronal sections of E17 brains stained for Tle4, a marker of LL neuronal identities. Scale bar of 20x objective lens: 50μm. (**H**) Quantification of neuronal positioning of Tle4 from pia to VZ. (**I**) Total amount of Tle4+ neurons as the percentage from the total *Gfp*+ neurons. *n*= 8-10 images from 2 Ctrl shRNA, *n*= 16-19 images from 3 animals for *DcpS* shRNA. Data represent the mean of sections and SEM. Statistics: Welch’s t-test per each bin, unpaired, 2-tailed or Mann-Whitney test. NS= not significant. *: p<0.05. VZ, Ventricular Zone. SVZ, Subventricular Zone. IZ, Intermediate Zone. LL, Lower-layer. UL, Upper-layer. MZ, Marginal Zone.

To analyze whether LL structures might be affected after DcpS KD, we also performed IHC against LL subcortically projecting Tle4 and Ctip2 expressing neurons. The laminar distribution of DcpS silenced Tle4+/Gfp+ and Tle4+/Satb2+/Gfp+ neurons (Fig. 5G and Fig S4A) showed modest increase of both in the LL (Fig. 5H and S4D). However, it did not lead to a significant increase in the total number of LL neurons (Fig. 5I and Fig. S4G). Lamination defects or change in the total neuronal amount were also not significant for LL Ctip2+/Brn1+/Gfp+ and Ctip2+/Gfp+ neurons between control and *DcpS* silenced cortices (Fig. S4B, E, H and Fig. S4C, F, I). Our results indicate that *DcpS* KD predominantly affects UL neuronal identities.

In addition, *DcpS* OE under the control of the *Cdk5r* promoter caused significant reduction of Satb2+ neurons in UL relative to the Ctrl OE (Fig. 6A and C). There was no apparent change in the neuronal distribution throughout other cortical zones. Also, *Cdk5r-DcpS* OE did not affect the positioning of the Brn1+/Gfp+ neurons (Fig. 6B and D). In addition, we observed no changes in the total Satb2+/Gfp+ or Brn1+/Gfp+ cell number relative to the control (Fig. 6E and F). Lastly, we assessed the consequences of *DcpS* OE on the LL specific neuronal subtypes. Surprisingly, the distribution of the Tle4+ neurons in the *Cdk5r-Dcps* OE (Fig. 6G) cortices was higher in the UL, LL and IZ (Fig. 6H). Moreover, *Cdk5r-DcpS* OE induced a significant increase in the total percentage of Tle4+/Gfp+ neurons relative to the Ctrl OE (Fig. 6I). The number of Satb2+/Tle4+/Gfp+ cells (Fig. S5C) was significantly upregulated in the UL and IZ when *DcpS* OE was driven by the *Cdk5r* promoter (Fig. S5F), which was associated with the increased total number of Satb2 Tle4+ neurons (Fig. S5I). In contrast, *Cdk5r-DcpS* OE did not alter the columnar distribution and the total amount of Ctip2+/Gfp+ and Ctip2+/Brn1+/Gfp+ neuronal subtypes (Fig. S5A, D, G and Fig. S5B, E, H). Therefeore, our observations suggest that changes seen upon *DcpS* OE are associated predominantly with Tle4+ neurons. Collectively, these data suggest that the scavenger enzyme DcpS plays an important role in regulating the laminar cell fate of neocortical neuronal subtypes. Specifically, our results show that DcpS regulates the balance in molecular identity of both UL and LL intracortically projecting neuronal subtypes and a subpopulation of LL subcortically projecting neurons.

**Figure 6.**
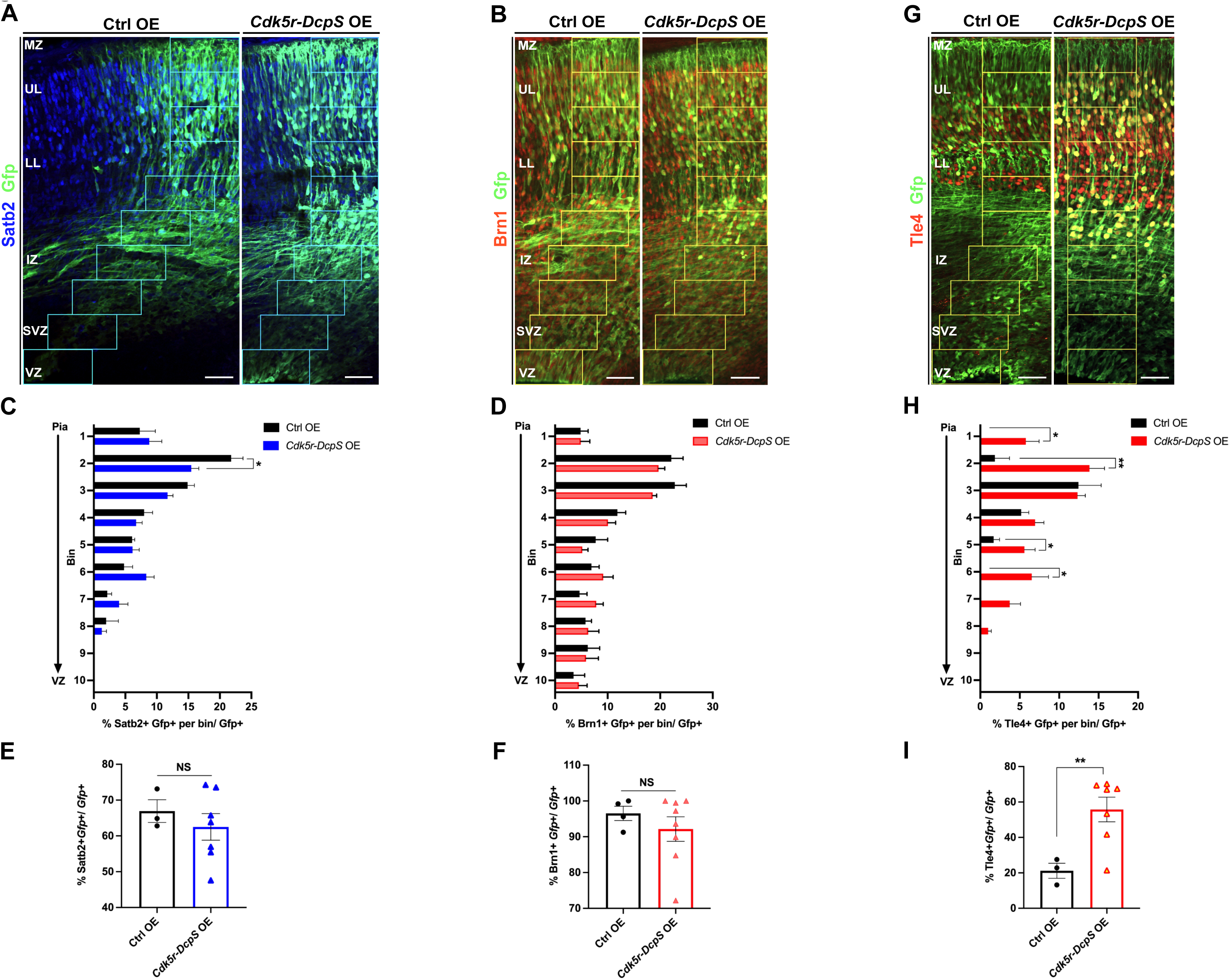
Overexpression of DcpS in post-mitotic neurons affects lower-layer neuronal subtype. Representative confocal images of E17 neocortices transfected at E13 with *CAG-GFP* and Ctrl OE or *Cdk5r-DcpS* OE construct. Neocortices were immunostained for upper layer identity markers (**A**) Satb2 in blue and (**B**) Brn1 in red. Scale bar of 20x objective lens: 50μm. (**C**) Quantified columnar distribution of Satb2+ and (**D**) Brn1+ neurons in each of the ten bins from pia to VZ as a percentage of the total *Gfp*+ cells from A and B. (**E**) Total number of *Gfp*+ Satb2+ or (**F**) *Gfp*+ Brn1+ neurons given as a percentage from total *Gfp+* cells. (**G**) Coronal sections of E17 brains immunostained for LL neuronal marker, Tle4+. Scale bar of 20x objective lens: 50μm. (**H**) Graphed *Gfp+Tle4+* cell fraction of the total *Gfp+* cells, quantified as described in C and D. (**I**) Total amount of Tle4+ neurons as the percentage from total *Gfp+* neurons. *n*= 4-8 images from 2 Ctrl OE, *n*= 14-15 images from 3 animals for animals for *Cdk5r-DcpS* OE. Data represent the mean of sections and SEM. Statistics: Welch’s t-test per each bin, unpaired, 2-tailed or Mann-Whitney test. NS= not significant. *: p<0.05, **: p<0.01. VZ, Ventricular Zone. SVZ, Subventricular Zone. IZ, Intermediate Zone. LL, Lower-layer. UL, Upper-layer. MZ, Marginal Zone.

## Discussion

Regulation of RNA metabolism has emerged as a critical modulator of neuronal development. We and others recently reported that the RNA stability associated DcpS scavenger decapping enzyme is essential for normal cognitive function (Ahmed I *et al.* 2015; Ng CK *et al.* 2015; Alesi V *et al.* 2018; Masoudi M *et al.* 2020). Here, we demonstrated a functional role for the scavenger decapping activity in determining neuronal development. iPSCs derived from individuals that only produce decapping deficient DcpS were compromised in their ability to differentiate into neurons, and to extend neuritic projections. In the mouse brain, manipulation of DcpS expression levels altered neuronal radial migration, axon growth and the subtype identity of neocortical neurons. Taken together, these results indicate an important role for this protein in neuronal development, which incorporate circuits important for complex behaviors. Specifically, DcpS was central for proper neuronal migration of upper layer neocortical neurons consistent with the higher order cognitive defects observed in individuals with defective DcpS decapping. Collectively, our studies reveal the contribution of DcpS, and more generally the significance of RNA metabolism, for normal development of neurons that contribute to the acquisition of complex behaviors, such as cognition.

Differentiation of patient derived iPSCs into excitatory glutamatergic iNs revealed that a functional scavenger decapping activity is necessary for the normal progression of neurogenesis. Patient iPSCs demonstrated diminished neuronal differentiation as determined by a reduction in the number of cells expressing the HuC/D neuronal marker (Fig. 1A and B) as well as an increased persistence in expression of the progenitor marker Pax6 in DcpS-Mut cells (Fig. 1C). These findings suggest that DcpS functions in the control of neuronal differentiation and neurogenesis. Furthermore, an important observation in these experiments was that patient cells generated fewer and shorter neuritic processes than control cells. Similarly, when endogenous *DcpS* mRNA was knocked down in mouse neurons these exhibited fewer and shorter callosal axonal projections (Fig. 3E), whereas neurons overexpressing DcpS displayed enhanced callosal axon outgrowth (Fig. 3F). Overexpression of DcpS in postmitotic migratory neurons not only promoted the growth of axons, but also favored the extension of apical dendrites (Fig. 3H and I). These findings suggest a critical role for DcpS in neural differentiation, neuritogenesis and migration through the regulation of a gene network that has yet to be determined.

The high level of DcpS expression in embryonic neocrtex (Fig. 2A and C) provided important insights into the precise subdomain of the brain where scavenger decapping activity may play a role. We hypothesized that neurons with reduced levels of DcpS may exhibit developmental abnormalities. Indeed, electroporation of *DcpS* shRNA in the embryonic mouse brain altered the laminar positioning of the radially migrating neurons in the neocortex (Fig. 3A). *DcpS* silencing hindered neuronal migration below their expected laminar positions (Fig. 3A and B). Selective overexpression of *DcpS* within postmitotic migratory neurons also affected radial migration, particularly that of upper-layer neurons (Fig. S2A, 3C and 3D). Furthermore, neurons electroporated with the *Cdk5r-Dcps* OE construct in postmitotic neurons also showed increased multipolar morphology, which may affect proper radial migration (Fig. 4B and C). Moreover, our GO-term functional annotation of the differentially expressed genes (Fig. 2E) showed that cellular components and biological processes were significantly reduced in *DcpS* KD mouse neurons and that they are mostly associated with the cell adhesion regulation and organization of dendrites and ECM.

Lastly, we investigated whether disruption of cortical migration following embryonic misexpression of DcpS was partially dependent on the targeted neuronal population. Our results indicated that knocking down endogenous *DcpS* levels severely affected the generation and distribution of the Brn1 and Satb2 neuronal subtypes (Fig. 5A-F), while having no effect on the LL. Conversely, overexpression of DcpS in post-mitotic neurons specifically impaired only the generation of LL Tle4+ cortical neurons (Fig. 6G-I) without influencing other neuronal identities. These observations propose a model in which reduced levels of DcpS impair laminar fates of UL intracortically projecting neurons, whereas higher levels of DcpS promote the production of the specific subpopulation of LL neurons. Thus, DcpS regulates the balance in UL and LL glutatamergic neurons, a phenomenon previously attributed to RNA control during neocrtical development (DeBoer EM *et al.* 2014; Popovitchenko T *et al.* 2020). Since our findings point to a critical and multipronged role for DcpS in the neocortical development, future studies should clarify the mechanism by which DcpS operates in postmitotic neurons to regulate the multiple steps of neuronal development.

The DcpS mutation studied in this report was originally identified within a consanguineous family in Pakistan (Ahmed et al. 2015) with individuals homozygous for a pre-mRNA splice site mutation (c.636+1G>A). This mutation alters an invariant G at the 5’ splice site of intron 4, leading to the utilization of a cryptic 5’ splice site 45 nucleotides downstream of the normal 5’ splice site. The aberrantly spliced mRNA encodes an in-frame 15 amino acid insertion (Val-Lys-Val-Ser-Gly-Trp-Asn-Val-Leu-Ile-Ser-Gly-His-Pro-Ala) after amino acid Q212 of the DCPS protein (DcpS-Mut). The mutation result in a loss of function phenotype with disrupted DcpS scavenger decapping activity (Ahmed I *et al.* 2015). As revealed in this study, leads to inefficient neurogenesis in iPSC neural differentiation and neurogenesis defects when knocked down in mouse neocortex.

An important direction for future study will be addressing how the scavenger decapping activity of DcpS modulates neural function. DcpS was initially identified as an activity that hydrolyzed the residual cap structure (m^7^GpppN) generated by the degradation of an mRNA from the 3’ end (Wang Z and M Kiledjian 2001; Liu H *et al.* 2002). One possibility is that in the absence of DcpS decapping, aberrant accumulation of m^7^GpppN may function as a signaling molecule that could modulate gene expression analogous to the function of numerous Nudix hydrolases that clear signaling dinucleotides (McLennan 2006). For example, cap structure accumulation could result in the sequestration of nuclear and cytoplasmic cap binding proteins to affect splicing and mRNA translation, respectively (Bail S and M Kiledjian 2008; Shen V et al. 2008). DcpS also contributes more directly to RNA stability (Zhou M et al. 2015). In the yeast Saccharomyces cerevisiae, the DcpS homolog, Dcs1 is an obligate cofactor for the 5’ to 3’ exoribonuclease, Xrn1 and necessary for 5’ end decay (Liu H and M Kiledjian 2005; Sinturel F et al. 2012) while in C. elegans, DcpS is involved in microRNA directed regulation through Xrn1, albeit through a decapping independent mechanism (Bosse GD et al. 2013; Meziane O et al. 2015). Moreover, a recent report demonstrated that the yeast Dcs1 protein also possesses canonical mRNA decapping function on m^7^G capped long RNA in addition to scavenger decapping on cap structure (Wulf MG et al. 2019) raising the prospect that DcpS may more directly alter the stability and accumulation of specific mRNAs. Future studies will address the molecular mechanism by which DcpS influences neurogenesis and human cognition.

## Acknowledgements

We thank Jennifer Moore and Michael Sheldon for their assistance with the generation of iPS cells.

---

**Supplementary Figure S1.**
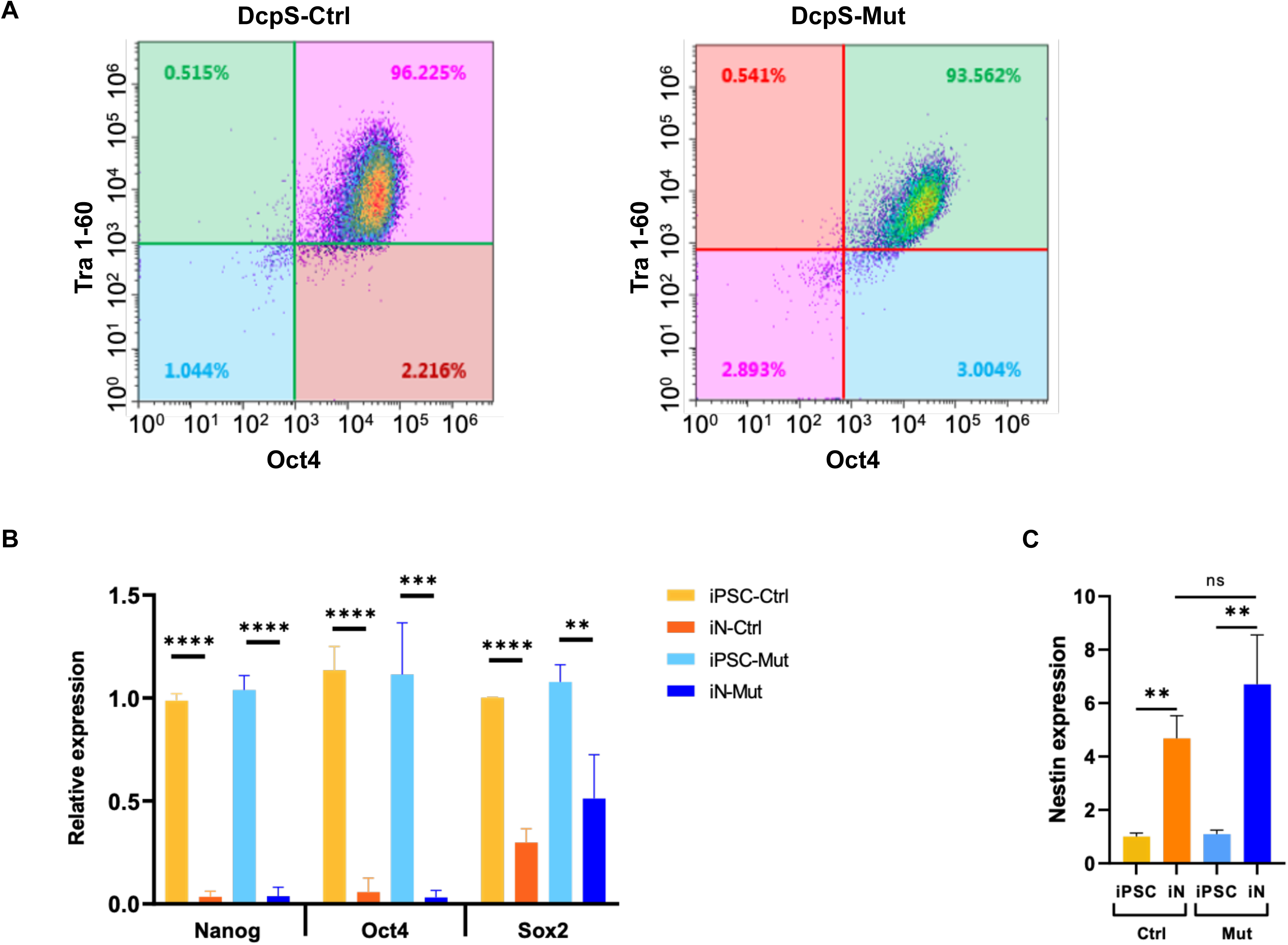
Expression of stem cell-specific markers in patient-derived iPSCs and iNs. (**A**) FACS analysis showing expression of stem cell markers Oct4/Tra-1-60. (**B**) Expression of Nanog, Oct4 and Sox2 in patient, iPSCs and Day7 iNs, detected using RT-qPCR. Error bars indicate standard deviation (SD). (**C**) Expression of Nestin as the marker for neuronal progenitor cells in Day 0 and Day 7 iNs, Error bars indicate standard error of mean (SEM).

**Supplementary Figure S2.**
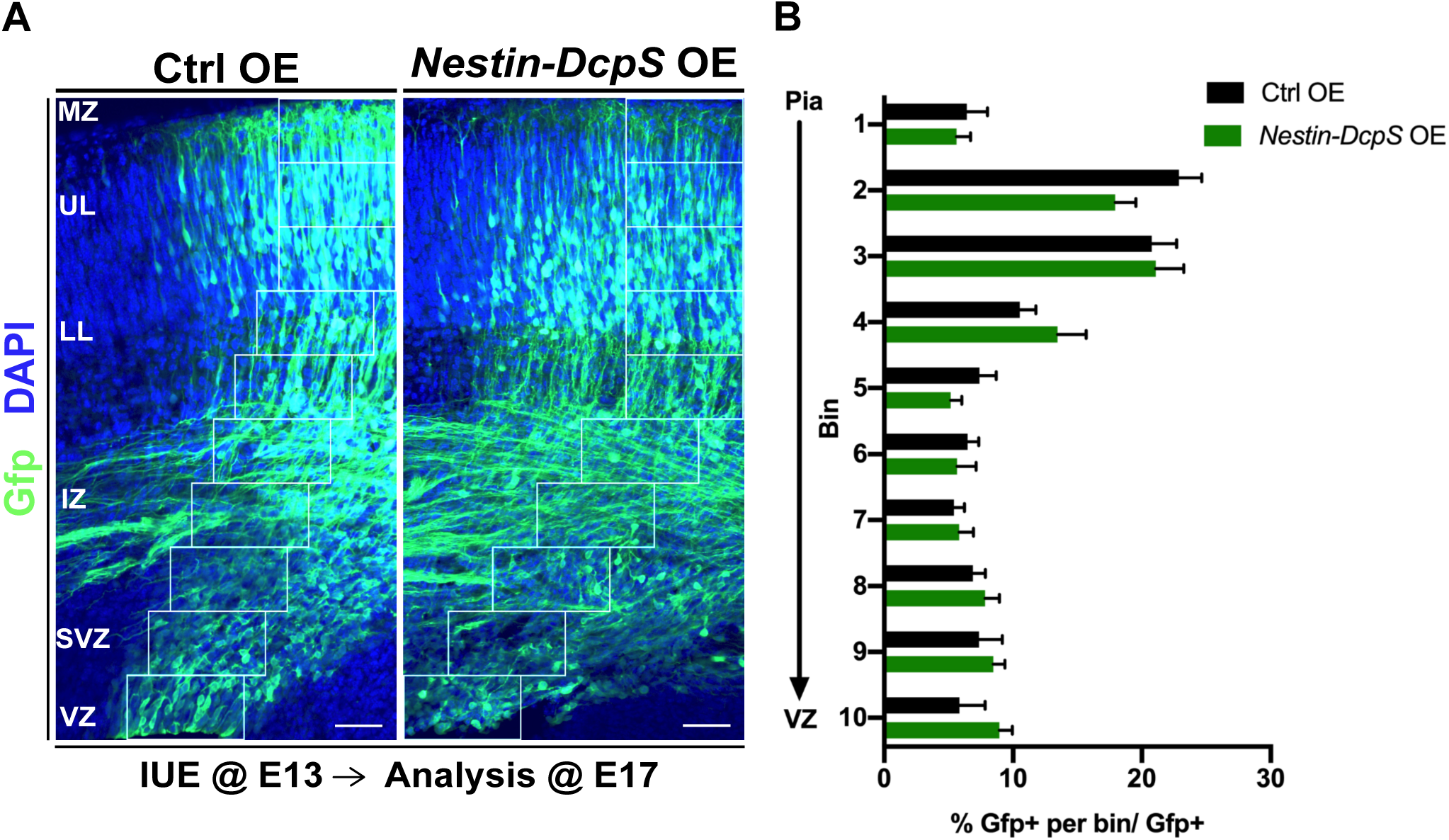
Minimal effect of DcpS on radial migration of early mitotic glutamatergic neocortical neurons. (**A**) Representative coronal confocal images of E17 neocortices transfected at E13 with *CAG-GFP* and Ctrl OE and *Nestin-DcpS* OE immunostained for *Gfp* (green) and DAPI (blue). Scale bar of 20x objective lens: 50μm. (**B**) Relative distribution of *Gfp+* cells subdivided in 10 equal bins for neuronal quantification from pial surface (bin 1) to VZ (bin 10) from A. Results represent *Gfp*+ neurons in each bin as the percentage from total *Gfp*+ neurons. *n*=12 images from 2 animals for Ctrl OE, *n*=20 images from 4 animals for *Nestin-DcpS*. Statistics: Welch’s t-test per each bin, unpaired, 2-tailed. *: p<0.05.

**Supplementary Figure S3.**
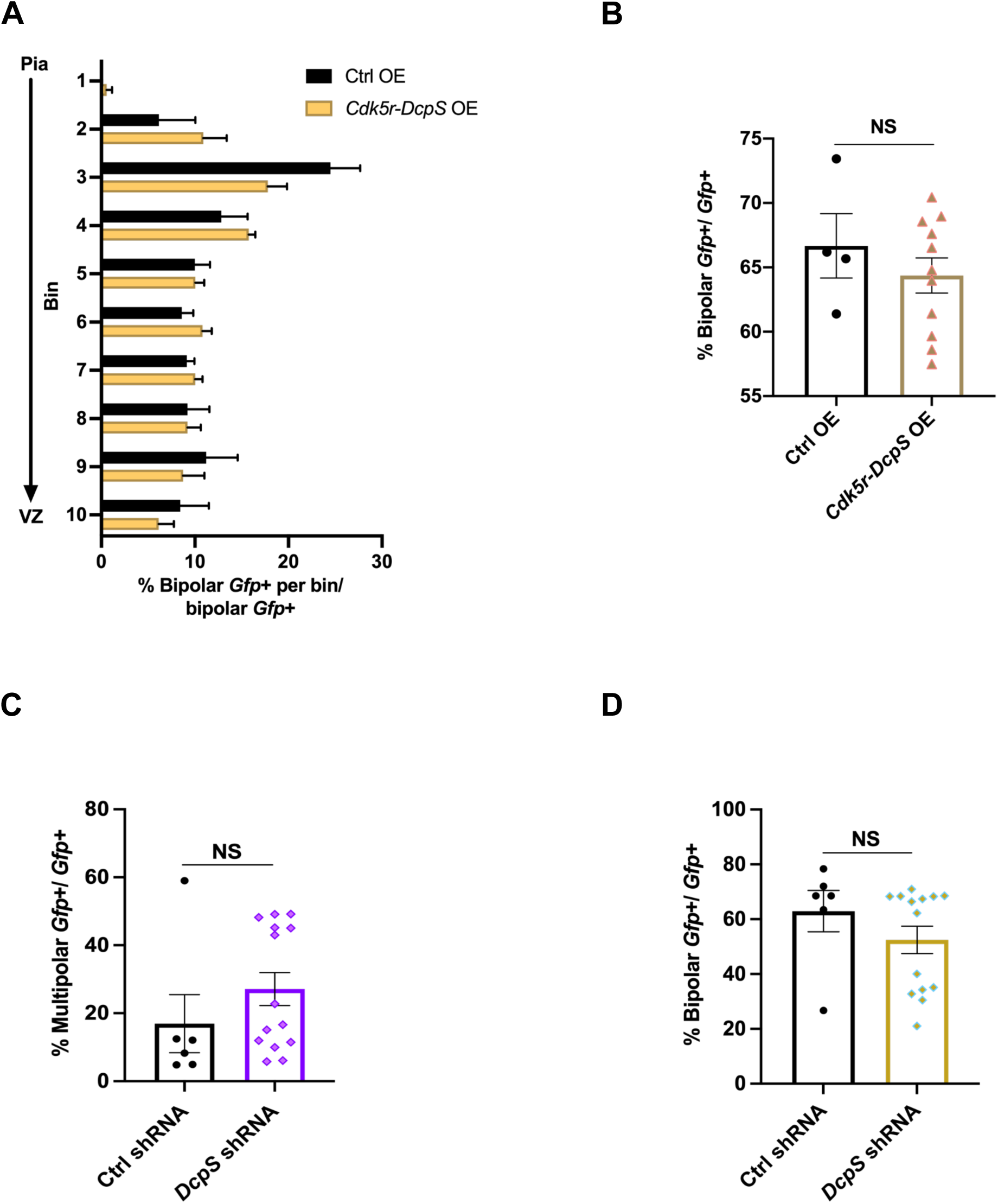
DcpS OE and knock down in postmitotic neurons. **A**) Distribution of bipolar *Gfp*+ neurons from Fig. 4A quantified as the percentage of bipolar *Gfp*+ neurons in each bin over the total morphologically specific *Gfp+* neurons. (**B**) Quantification of the total proportion of bipolar *Gfp+* neurons transfected with Ctrl or *Cdk5r-DcpS* OE from the total *Gfp+* labeled cells. *n=* 4 images from 2 animals for Ctrl OE, *n=* 11 images from 3 animals for *Cdk5r-DcpS* OE condition. (**C**) Total multipolar *Gfp*+ neurons, or (**D**) bipolar *Gfp*+ neurons quantified as a proportion of total *Gfp+* neurons. *n*= 6 images from 2 animals for Ctrl shRNA, *n*= 14 images from 3 animals for DcpS shRNA. Data represent the mean of sections and SEM. Statistics: Welch’s t-test per each bin, unpaired, 2-tailed, relative to the control. NS: not significant.

**Supplementary Figure S4.**
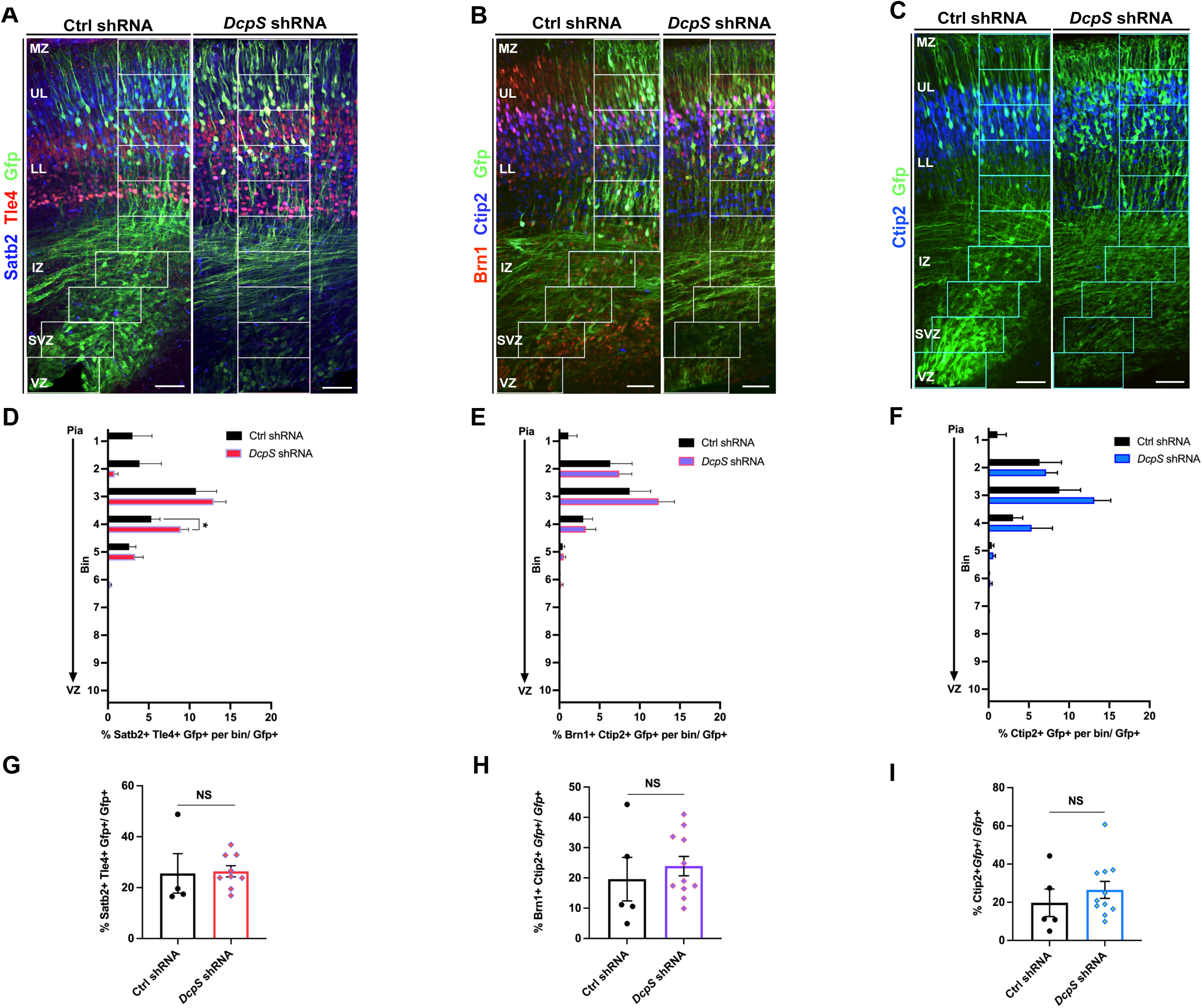
Quantification of subpopulation of glutamatergic neurons in *DcpS* knockdown conditions. E13 embryos were electroporated with Ctrl shRNA or *DcpS* shRNA constructs together with *CAG-GFP*. (**A**) Coronal sections of E17 brains co-stained for Satb2 and Tle4, (**B**) Brn1 and Ctip2, or (**C**) Ctip2, established markers of either UL or LL neurons. Scale bar of 20x objective lens: 50μm. (**D**) Laminar distribution of *Gfp*+ cells that coexpressed Satb2 and Tle4, (**E**) Brn1 and Ctip2, or (**F**) solely Ctip2 presented as fraction of total *Gfp+* cells. (**G**) Quantification of Satb2+ Tle4+, (**H**) Brn1+ Ctip2+, or (**I**) Ctip2+ neurons presented as a total of *Gfp+* cells. *n*= 8-10 images from 2 Ctrl shRNA, *n*= 16-19 images from 3 animals for *DcpS* shRNA. Data represent the mean of sections and SEM. Statistics: Welch’s t-test per each bin, unpaired, 2-tailed or Mann-Whitney test. NS= not significant. *: p<0.05. VZ, Ventricular Zone. SVZ, Subventricular Zone. IZ, Intermediate Zone. LL, Lower-layer. UL, Upper-layer. MZ, Marginal Zone.

**Supplementary Figure S5.**
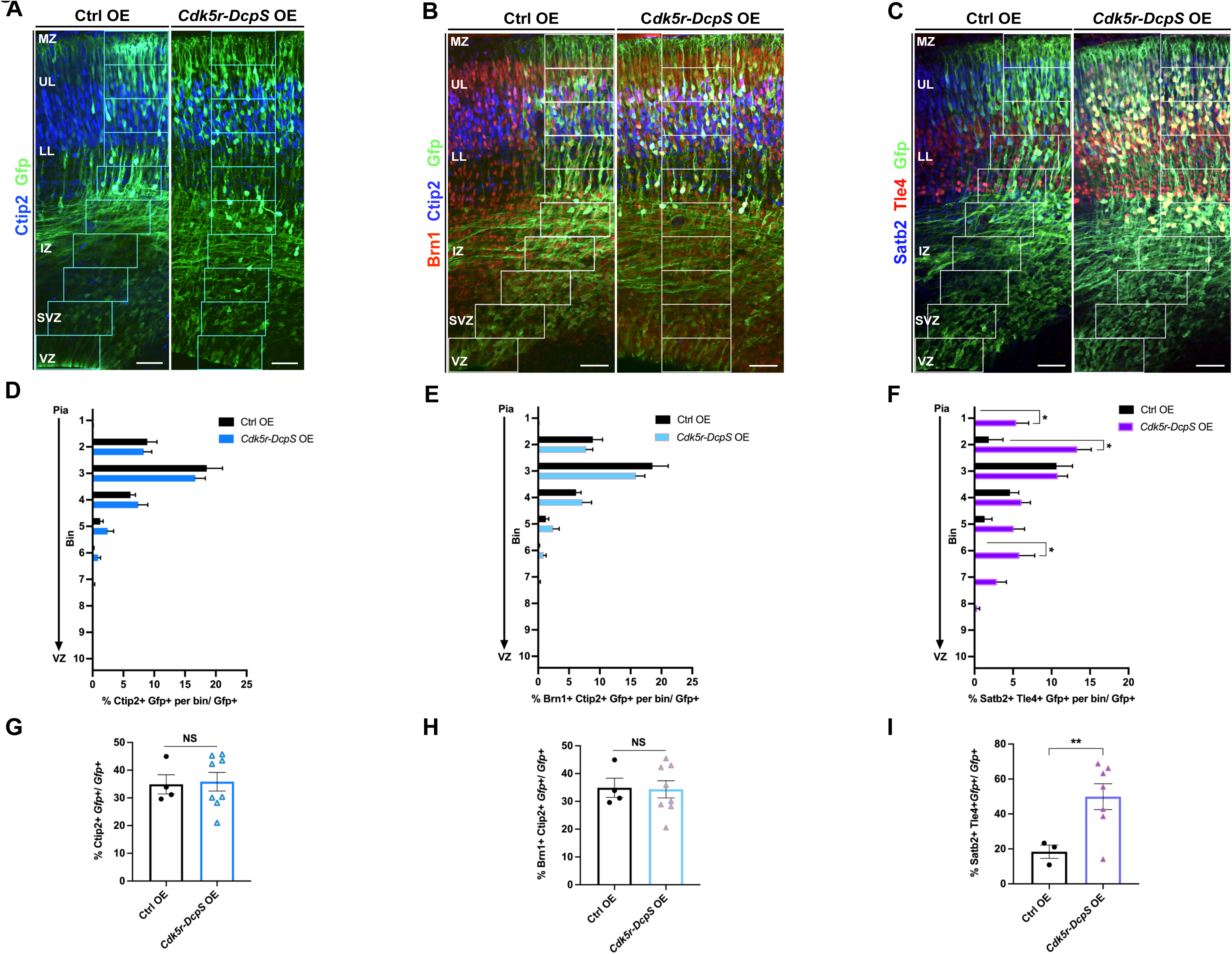
DcpS overexpression in postmitotic neurons increases the total number of co-localized Satb2+ and Tle4+ neurons. (**A**) Coronal sections of E17 brains electroporated with Ctrl OE or Cdk5r-DcpS OE constructs at E13 and stained against Ctip2, (**B**) Brn1 and Ctip2, and (**C**) Satb2 and Tle4. Scale bar of 20x objective lens: 50μm. (**D-F**) Quantification of specific neuronal identities in each of the ten bins and (**G-I**) in total as a percentage from all *Gfp+* cells. *n*= 4-8 images from 2 Ctrl OE, *n*= 14-15 images from 3 animals for *Cdk5r-DcpS* OE. Data represent the mean of sections and SEM. Statistics: Welch’s t-test per each bin, unpaired, 2-tailed or Mann-Whitney test. NS= not significant. *: p<0.05, **: p<0.01.

